# *In vitro* nonalcoholic fatty liver disease model with cyclo-olefin-polymer-based microphysiological systems

**DOI:** 10.1101/2020.12.28.424535

**Authors:** Xiaopeng Wen, Makoto Yamanaka, Shiho Terada, Ken-ichiro Kamei

## Abstract

Nonalcoholic fatty liver disease (NAFLD) is one of the common chronic liver conditions, whose treatment involves curing patients without liver transplantation. Understanding the mechanism of NAFLD initiation and progression would enable development of new diagnostic tools and drugs; however, until now, the underlying mechanisms of this condition remain largely unknown owing to the lack of experimental settings that can simplify the complicated NAFLD process *in vitro*. Microphysiological systems (MPSs) have long been used to recapture human pathophysiological conditions *in vitro* for applications in drug discovery. However, polydimethylsiloxane (PDMS) has been used in most of these MPSs as the structural material; it absorbs hydrophobic molecules, such as free fatty acids (FFAs), which are the key components that initiate NAFLD. Therefore, the current PDMS-based MPSs cannot be directly applied to *in vitro* NALFD modeling. In this work, we present an *in vitro* NAFLD model with an MPS made of cyclo-olefin polymer (COP), namely COP-MPS, to prevent absorption of FFAs. We demonstrated induction of the NAFLD-like phenotype in HepaRG hepatocyte-like cells cultured in the COP-MPS by introducing FFAs. The FFAs induced lipid accumulation in the HepaRG cells, resulting in inactivation of the apoptotic cells. We believe that the proposed COP-MPS can contribute toward investigations of NAFLD mechanisms and identification of new drugs to prevent the progression of liver disease and avoid liver transplantation.

## 1. Introduction

Nonalcoholic fatty liver disease (NAFLD) is a common chronic liver condition associated with metabolic syndrome, such as obesity, hypertension, hyperlipidemia, cardiovascular dysfunction, systemic inflammation, and many extrahepatic diseases (Byrne and Targher, 2015; Younossi et al., 2019; Younossi, 2019). Until now, liver transplantation is still the only means of preventing the progression of and curing liver diseases, but it is extremely difficult to match donors to patients. Therefore, there is an urgent need to find alternative means to cure patients, with NAFLD being one of the critical stages in the progression of liver diseases. However, NAFLD is a complex disease that encompasses a broad spectrum of hepatic phenotypes, including nonalcoholic steatohepatitis (NASH), liver fibrosis, and inflammation (Tilg et al., 2020), and the underlying mechanisms of NAFLD progression are still largely unknown. Thus, there are no effective treatments and drugs to cure patients with NAFLD.

To elucidate the mechanisms of NALFD progression, experimental models, such as animal models (Liu et al., 2013) and *in vitro* human models (Boeckmans et al., 2018; Davidson et al., 2016), have been used in literature. On the one hand, animal models allow addressing the mechanism of NAFLD progression *in vivo*; however, species differences cannot be ignored. On the other hand, *in vitro* human models using human primary hepatocytes or cell lines have been used to simplify NAFLD for better understanding and application to drug screening. Further, most cell-culture methods that are currently used involve cell-culture plates and flasks, which cannot represent the *in vivo* physiological conditions, thus resulting in less functionality of the cultured hepatocytes.

To address these issues, microphysiological systems (MPSs) or organ/body-on-a-chip (O/BoC) systems based on microfluidic technology offer the possibility of simulating human physiological and pathological conditions *in vitro* for application in drug development, toxicological tests, and disease modeling (Benam et al., 2015; Kamei et al., 2017; Kimura et al., 2018; Soto-Gutierrez et al., 2017; Sung et al., 2019). MPS platforms provide dynamic and three-dimensional culture conditions that are unavailable in conventional cell culture methods to improve cellular functions. However, MPS platforms still have some issues to be addressed, particularly those based on polydimethylsiloxane (PDMS), which has often been used in the fabrication of MPS platforms owing to its biocompatibility, gas permeability, and transparency. Since PDMS also exhibits hydrophobicity and porosity, which allows chemical/drug absorption, water evaporation, deformation, and leaching of un-crosslinked reagents (Berthier et al., 2012; Carter et al., 2020; Toepke and Beebe, 2006; van Meer et al., 2017), PDMS-based MPSs have limited applications owing to the undesired influence of the cellular responses (Kamei et al., 2013). In particular, to establish *in vitro* NAFLD models, the absorption of hydrophobic molecules is considered a critical property because the key molecules for NAFLD modeling, such as free fatty acids (FFAs), are absorbed into the PDMS material to reduce the amount of FFAs in the cell-culture media. Moreover, PDMS absorption also reduces the tested drugs/chemicals in cell culture media, thereby causing misleading results on drug efficacy and toxicity. Therefore, there is a need for alternative materials for *in vitro* NAFLD modeling.

To fulfill these requirements, cyclo-olefin polymer (COP) has been proposed as a suitable material over PDMS because of its chemical/physical stability, high purity, and optical clarity (Campbell et al., 2020; Illa et al., 2010). Moreover, COP materials have thermostability and high modulus for metal molding processes and can be applied to mass produce MPSs. Although the previously reported microfluidic devices made of COP require solvents or adhesive tapes to assemble the COP materials, which may affect cellular phenotypes (Wan et al., 2017), the photobonding process using vacuum ultraviolet (VUV) from an excimer light source allows fabrication of the COP-based MPS without the use of additional solvents and adhesives (Yamanaka et al., 2020).

Herein, we introduce an *in vitro* NAFLD model with an MPS platform made of COP. We confirmed that the COP-MPS has the ability to prevent absorption of hydrophobic molecules, unlike PDMS-MPS. Therefore, after FFA treatment for hepatocyte-like cells (HLCs), we found that the cells cultured in the COP-MPS could be observed with AdipoRed fluorescent cellular lipid staining, whereas the HLCs in the PDMS-MPS could not be observed owing to the strong background signals of the AdipoRed absorbed in the PDMS material. Furthermore, HLCs in the COP-MPS show intracellular lipid accumulation, resulting in induction of apoptosis. Thus, COP-MPS promises to improve the MPS for applications to *in vitro* NAFLD modeling for investigating the mechanisms behind NAFLD initiation and progression and aiding drug discovery.

## 2. Materials and Methods

### 2.1 Fabrication of the COP-MPS

The COP-MPS comprises two layers of COP and eight microfluidic cell-culture channels (800 μm width, 7500 μm length, 250 μm height), with a medium inlet and outlet each (2 and 1 mm in diameter, respectively) and a total channel volume of 13.2 μL each (Yamanaka et al., 2020). The methods used to fabricate COP-MPS include metal molding (Fig. S1A) followed by photobonding (Fig. S1B), as described previously (Yamanaka et al., 2020). Briefly, two metal molds were used to fabricate the microfluidic structures. The COP material (Zeonex 480R, Zeon, Tokyo, Japan) was injected into the two molds individually. After removing the structured COP components from the molds, the components were irradiated with VUV from an excimer lamp source (172 nm; Ushio INC., Tokyo, Japan) at 25 °C. The component surfaces were then assembled using a heat press at < 132 °C. Finally, ethylene oxide gas (Japan Gas Co. Ltd., Kanagawa, Japan) was used to decontaminate the fabricated COP-MPS.

### 2.2 Fabrication of the PDMS-MPS

Polymethyl methacrylate (PMMA, Sumitomo Chemical Co., Ltd., Tokyo, Japan) was used along with the micromilling procedure (CNC JIGBORER YBM 950V, Yasda Precision Tools K. K, Okayama, Japan) to fabricate the mold for the PDMS-MPS. Pre-PDMS (Sylgard184; A and B) was mixed at a 10:1 ratio and degassed for 1 h at 25 °C. The mixed pre-PDMS was then poured over the PMMA mold and placed in an oven at 65 °C for 2 h (Fig. S2). The solidified PDMS was then peeled off the mold and placed in an oven at 80 °C for 24 h. The surface of the PDMS was cleaned by soaking in ethanol (EtOH) and sonication. The PDMS part and a glass slide (Matsunami Glass, Osaka, Japan) were subsequently treated with VUV from an excimer lamp source (172 nm, Ushio INC.) at 25 °C and assembled at 80 °C for 2 days. Before being used for cell culture, this PDMS device was irradiated with UV radiation for decontamination.

### 2.3 HepaRG cell culture in MPS

Prior to cell culture in the MPSs, about 15 μL of 0.1% (w/v) gelatin (bovine serum albumin: BSA, Sigma-Aldrich, St. Louis, MO, USA) in distilled water was introduced into each microfluidic cell culture channel and maintained at 4 °C for over 24 h. The excess gelatin was then removed, and the coated channels were washed with fresh HepaRG medium 670 (KAC Co., Ltd., Tokyo, Japan). Differentiated HepaRG cells were obtained from KAC Co., Ltd., and the cells were maintained in the HepaRG medium 670. Prior to introduction in the MPSs, the cells were cultured in a humidified incubator at 37 °C with 5% (v/v) CO_2_. The cells were then harvested with trypsin/EDTA (0.04%/0.03%[v/v]) solution. After centrifugation, the cells were re-suspended in fresh cell-culture medium, and about 15 μL of 2.0 × 10^6^ cells mL^-1^ of the cell suspension was introduced into each microfluidic cell-culture chamber.

### 2.4 Free fatty acid treatment

Palmitic acid sodium salt (PA, Nakalai Tesque, Kyoto, Japan) and oleic acid sodium salt (OA, Nakalai Tesque) were used as the FFAs in this study. The PA and OA stock solutions were individually prepared by dissolving in phosphate-buffered saline (PBS) containing 30% (w/v) BSA (Sigma-Aldrich) solution at 50 mM for each sample. The OA:PA mixed solution was prepared at a 1:2 molecular ratio (OA 33.33 mM and PA 16.67 mM), and this solution was diluted in the HepaRG medium to obtain a total FFA concentration of 500 μM as the working solution. As a negative control, 30% (w/v) BSA in PBS was used. The platform was then incubated at 37 °C in a humidified incubator for 24 h, with the medium being exchanged every 12 h.

### 2.5 Staining for intracellular lipid accumulation

Lipid accumulation was visualized with an AdipoRed assay (Lonza, Basel, Switzerland) according to the manufacturer’s protocol. Briefly, the cells were fixed with 4% (v/v) PFA in PBS for 20 min at 25 °C. Next, about 15 μL of the AdipoRed assay reagent was mixed with 1 mL of PBS to obtain the AdipoRed staining solution. Then, 10 μL of this AdipoRed staining solution was introduced and incubated at 37 °C for 15 min. The chambers were washed with PBS three times and lastly with 300 nM of DAPI at 25 °C for 30 min.

### 2.6 Staining for apoptotic cells

Annexin V staining was performed according to the product manual of Alexa Fluor 594-Annexin V conjugate (Molecular Probes, Eugene, OR). Briefly, after washing with annexin-binding buffer (10 mM HEPES, pH 7.4, 140 mM NaCl, 2.5 mM CaCl_2_), the cells were stained with the Alexa Fluor 594-Annexin V conjugate at 25 °C for 15 min. Following cell fixation with 4% (v/v) PFA in PBS at 25 °C for 15 min, the cells were incubated with 300 nM Hoechst 33258 at 25 °C for 30 min.

### 2.7 Microscopic cell imaging

The samples containing cells were placed on the imaging stage of a Nikon ECLIPSE Ti inverted fluorescence microscope equipped with a CFI plan fluor 10×/0.30 NA objective lens (Nikon, Tokyo, Japan), CCD camera (ORCA-R2; Hamamatsu Photonics, Hamamatsu City, Japan), mercury lamp (Intensilight; Nikon), XYZ automated stage (Ti-S-ER motorized stage with encoders; Nikon), and filter cubes for the fluorescence channels (DAPI, GFP, HYQ, and TRITC; Nikon).

### 2.8 Single-cell profiling based on microscopic images

Following the microscopy image acquisition, CellProfiler software (Broad Institute of Harvard and MIT, Version 3.1.9) was used to identify the cells using Otsu’s method (McQuin et al., 2018). The fluorescence signals of the individual cells were quantified automatically. Violin and box plots were then generated using R software (ver. 3.5.2; https://www.r-project.org/).

### 2.9 Statistical analysis

The P values were estimated by the Steel–Dwass test using R software (ver. 3.5.2; https://www.r-project.org/).

## 3. Results

### 3.1 HepaRG cell culture in COP-MPS

One of the early steps in NAFLD progression in the liver is lipid accumulation after being delivered from the intestine. Therefore, treatment with FFAs for hepatocytes is commonly used as an *in vitro* NAFLD model in general cell culture. However, the MPS platforms made of PDMS have critical issues with absorption of hydrophobic molecules, including FFAs. In this study, we fabricated an MPS made of COP to prevent absorption of the FFAs for better establishment of an *in vitro* NAFLD model in the MPS (COP-MPS), as shown in Fig. 1A, 1B, and S1. This COP-MPS has eight microfluidic cell-culture chambers with medium inlet and outlet (Fig. 1C). For comparison, an MPS made of PDMS with the same microfluidic structure was also fabricated (Fig. S2).

**Fig. 1.**
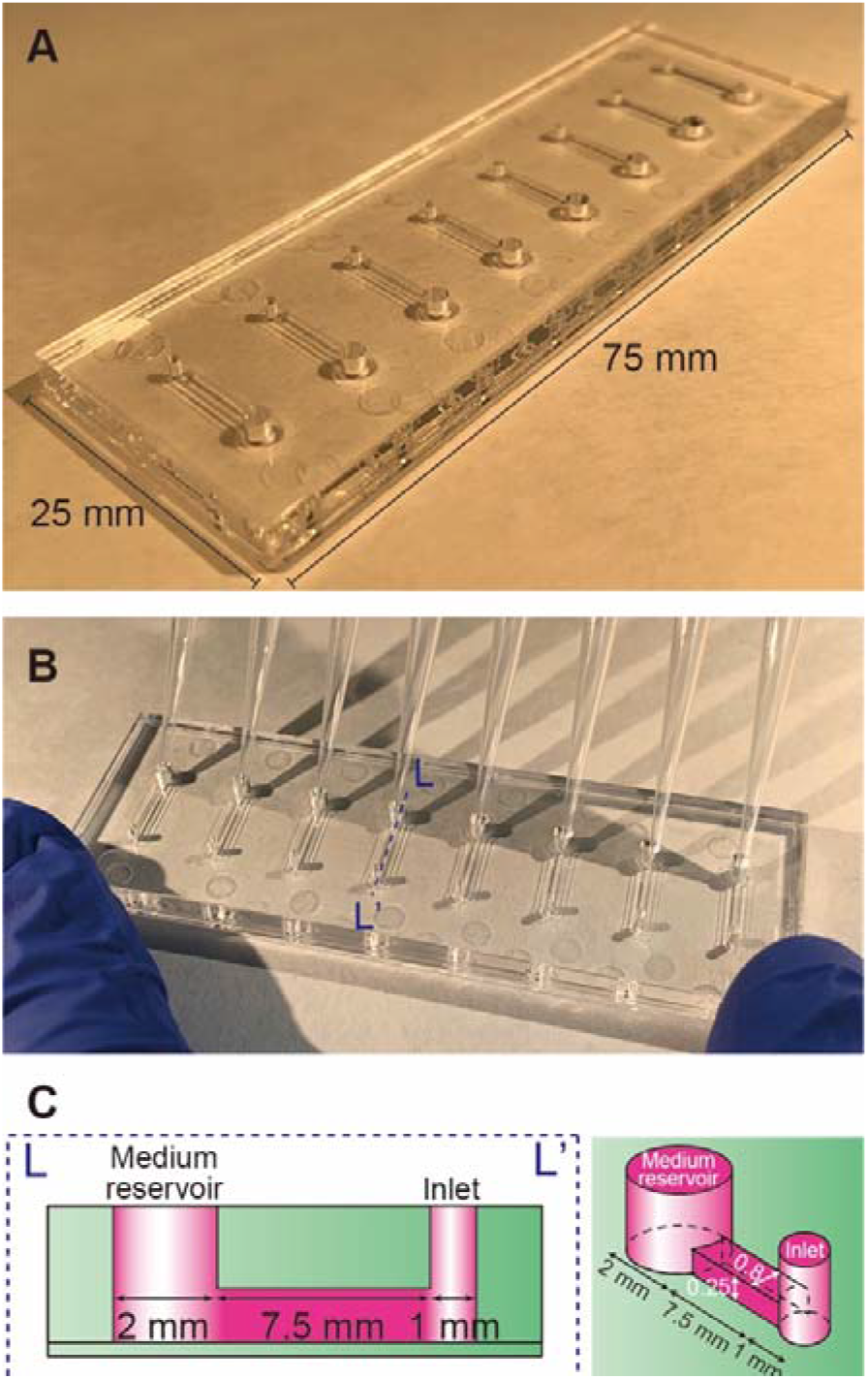
(A) Photograph of a microphysiological systems (MPS) made of cyclo-olefin polymer (COP), namely COP-MPS, for modeling nonalcoholic fatty liver disease (NAFLD). (B) Photograph of COP-MPS with eight tips and an octa-pipette to introduce cells and solutions into the microfluidic cell culture chambers simultaneously. (C) Schematic representation of a microfluidic cell culture chamber (L-L’) of the COP-MPS shown in (B).

In terms of the hepatocytes, HepaRG cells were used in this study for their ability to proliferate at the undifferentiated stage and differentiate into functional HLCs. Differentiated HepaRG cells have also been used as cell sources for *in vitro* NAFLD models in literature (Rogue et al., 2014).

### 3.2 Treatment of FFAs induce lipid accumulation in HepaRG cells cultured in COP-MPS

Prior to establishing the *in vitro* NAFLD model in the MPS, differentiated HepaRG cells were introduced in both the COP- and PDMS-MPS. To induce NAFLD-like phenotypes *in vitro*, a mixture of FFAs (palmitic acid (PA; 16:0) and oleic acid (OA; 18:1), which are the most abundant in both healthy subjects and patients with NAFLD) was used (Gómez-Lechón et al., 2007; Yao et al., 2011), and BSA was used as the carrier for FFAs in the cell-culture medium. Prior to FFA treatment, the absorption of hydrophobic molecules into the materials used for the MPS was evaluated using the AdipoRed fluorescent compound (Fig. 2). After 12 h of incubation of the MPS microfluidic channels with AdipoRed, the excess AdipoRed was rinsed with fresh cell-culture medium. A strong AdipoRed fluorescence signal was observed in the PDMS-MPS, while the COP-PDMS did not demonstrate any fluorescence (Fig. 2A). In the case of the PDMS-MPS, an AdipoRed fluorescence signal was observed not only in the microfluidic channel but also in the PDMS material, suggesting that the AdipoRed not only covered the PDMS surface but was also absorbed into the material. According to the quantitative results based on the microscopy images shown in Fig. 2A, the COP-MPS showed very low intensity fluorescence signals within the microfluidic channel, while the PDMS-MPS showed 10.4-fold stronger fluorescence signals (Fig. 2B).

**Fig. 2.**
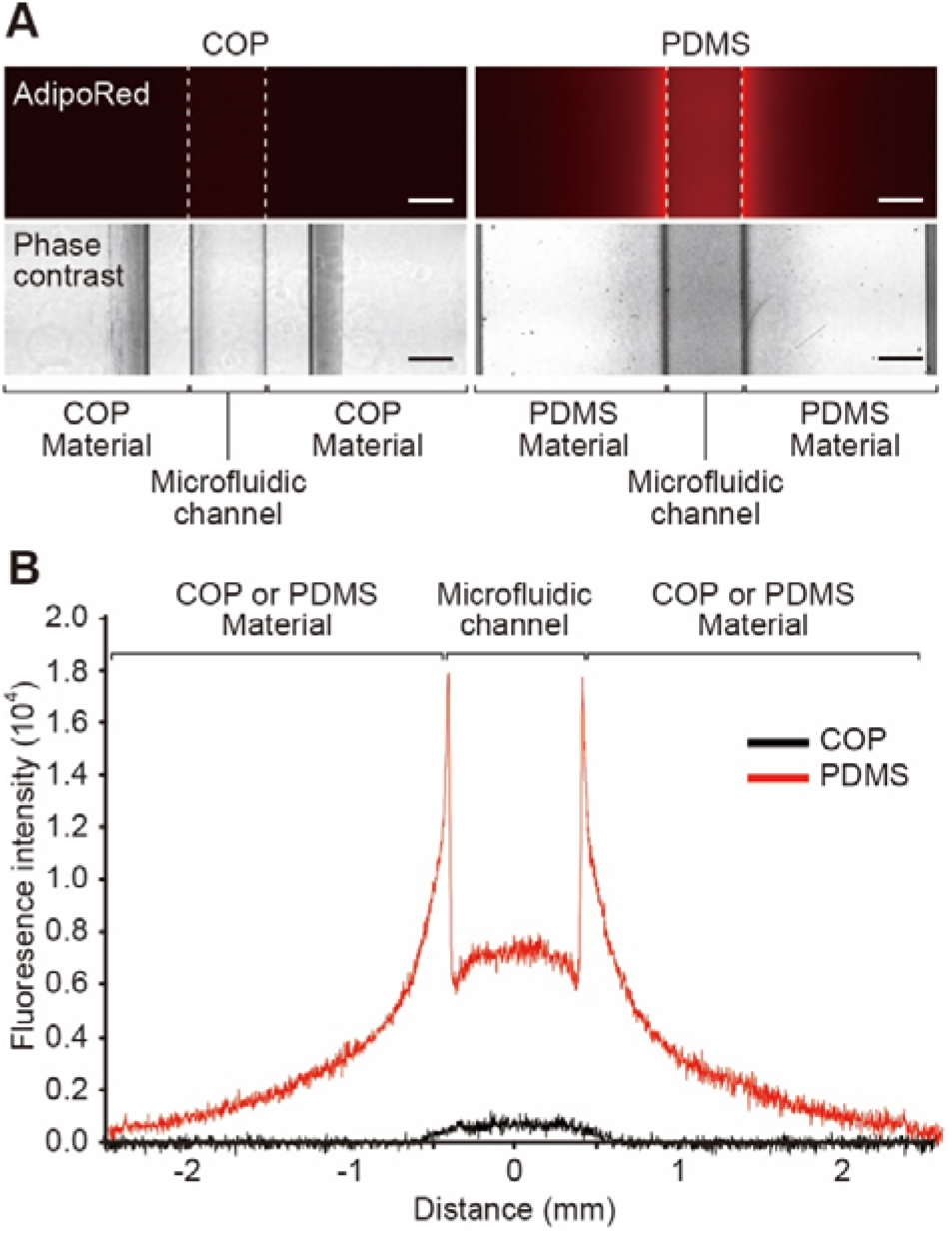
Evaluation of molecular absorption into the COP and PDMS used in the MPS. (A) Micrographs of the COP- and PDMS-MPS microfluidic channels treated with AdipoRed fluorescent compound (excitation: 553 nm; emission: 637 nm) for 12 h at 25 °C. The scale bars represent 500 μm. (B) Fluorescence intensity profiles across the COP- and PDMS-MPS microfluidic channels in (A).

Differentiated HepaRG cells in the COP- and PDMS-MPS were treated with FFAs and stained with AdipoRed to visualize lipid accumulation in the HepaRG cells (Fig. 3). Since the PDMS-MPS absorbed the FFAs as well as AdipoRed, the cells could not be observed clearly owing to the strong background fluorescence from the PDMS-MPS (Fig. 3A). Conversely, cellular lipid accumulation was observed for the COP-MPS (Fig. 3B). Therefore, the PDMS-MPS could not be used for further analyses, and only the COP-MPS was used. Treatment with FFAs (PA and OA:PA) and BSA-induced lipid accumulation in differentiated HepaRG cells and PA treatments showed the highest accumulations in the tested samples.

**Fig. 3.**
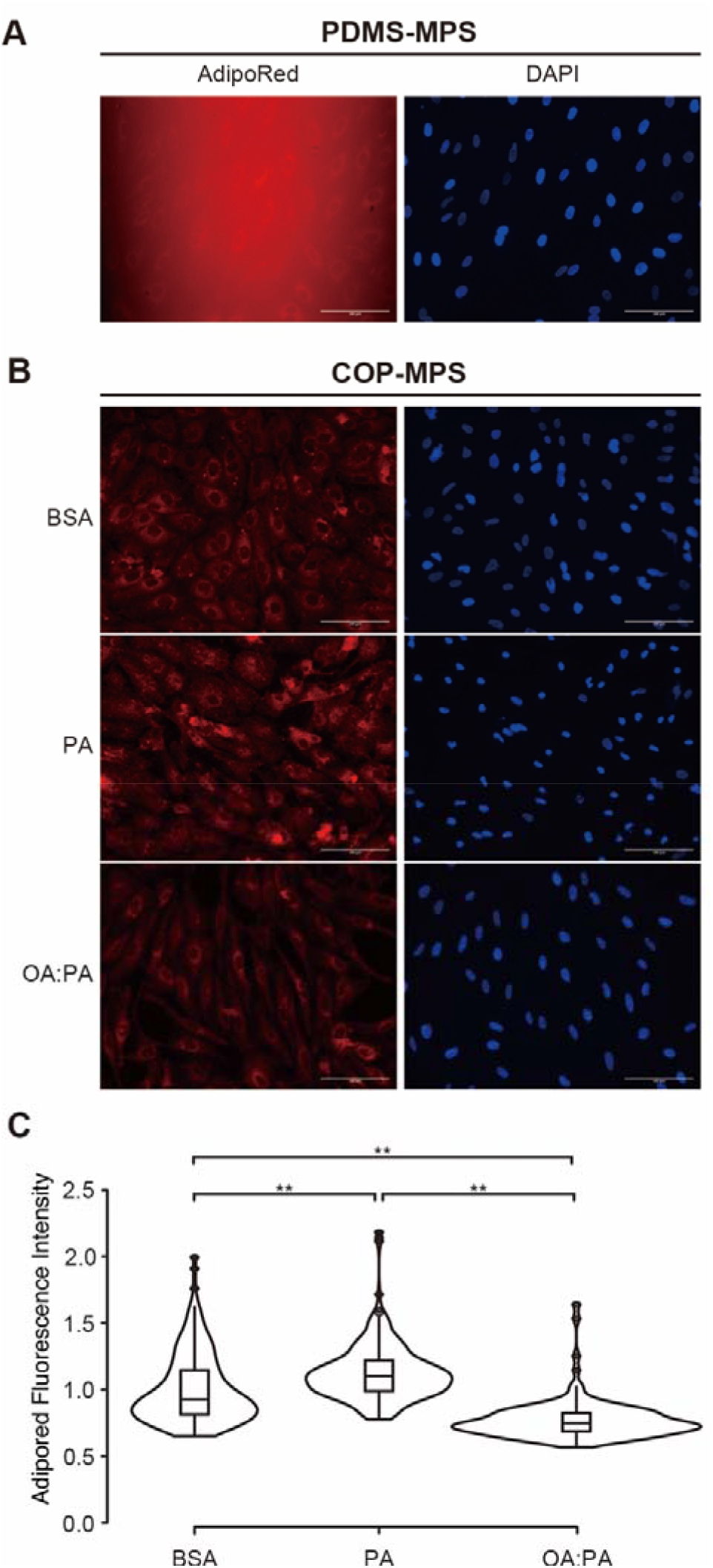
Observation of lipid accumulation in HepaRG cells by AdipoRed fluorescent lipid marker. (A) Fluorescence micrographs of the HepaRG cells cultured in PDMS-MPS treated with AdipoRed dye. DAPI was used to visualize the cellular nuclei. The scale bars represent 100 μm. (B) Fluorescence micrographs of the HepaRG cells cultured in COP-MPS treated with AdipoRed dye. The scale bars represent 100 μm. (C) Violin plots of single-cell profiles of AdipoRed fluorescence intensities of the individual FFA-treated HepaRG cells. \*\**p* < 0.01.

### 3.3 Accumulated lipids cause apoptosis of HepaRG cells cultured in COP-MPS

To evaluate whether the accumulated lipids induced apoptosis, cellular staining with Annexin V apoptotic marker was conducted for the FFA-treated HepaRG cells in COP-MPS (Fig. 4). Regarding the fluorescence micrographs of the Annexin V stained and quantitative single-cell profiles, PA measurements for 24 h showed maximal apoptotic cells compared to other treatments, such as those with BSA and OA:PA (Fig. 4A and B). Moreover, the average nuclear size of the HepaRG cells was evaluated since the apoptotic cells showed shrinkage of the nuclei (Fig. 4C). While the BSA- and OA:PA-treated HepaRG cells showed 656.2 ± 18.0 and 647.8 ± 11.1 μm^2^, respectively, the PA-treated cells showed 553.5 ± 18.6 μm^2^, showing nuclear shrinkage due to apoptosis. These results are in good agreement with the cellular lipid accumulations shown in Fig. 3, suggesting that lipid accumulation causes cellular apoptosis in HLCs.

**Fig. 4.**
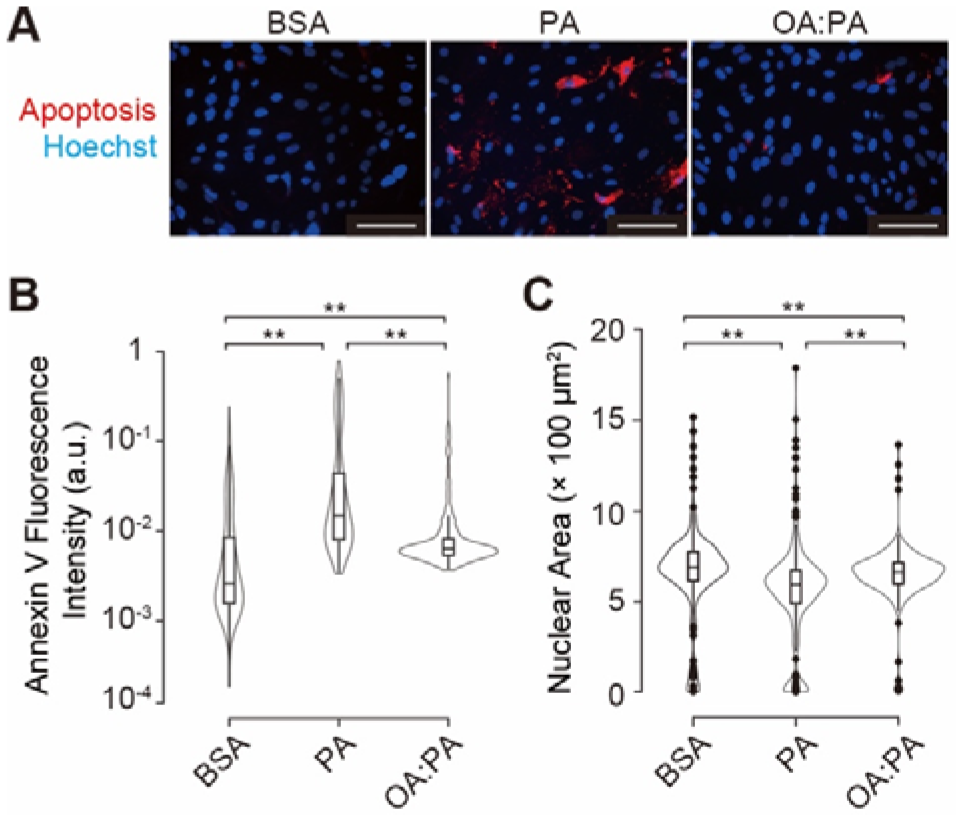
Observation of apoptosis in HepaRG cells by Annexin V fluorescent apoptosis marker. (A) Fluorescence micrographs of the HepaRG cells cultured in PDMS-MPS treated with Annexin V. Hoechst 33258 was used to visualize the cellular nuclei. The scale bars represent 100 μm. (B) Violin plots of single-cell profiles of Annexin V fluorescence intensities of individual FFA-treated HepaRG cells. \*\**p* < 0.01. (C) Violin plots of single-cell profiles of nuclear sizes of individual FFA-treated HepaRG cells. \*\**p* < 0.01.

## 4. Discussion

PDMS has been widely used in the fabrication of microfluidic devices, including MPS. However, PDMS usage has been associated with problems in cell culture and applications in drug development and disease modeling (Berthier et al., 2012; Carter et al., 2020; Toepke and Beebe, 2006; van Meer et al., 2017). Among these problems, absorption of the hydrophobic molecules is critical for drug development and disease modeling. Many drugs and their candidates are based on organic compounds and often have aromatic or similar structures, showing hydrophobicity. In addition, to establish an *in vitro* disease model, both genetic and chemical approaches are used, where some chemical approaches require hydrophobic molecules to recapture the pathological cellular status, e.g., FFAs for NAFLD modeling. MPS with the conventional PDMS material for NAFLD modeling has been traditionally limited by the aforementioned difficulty; thus, an alternative material is required for the MPS.

To date, there have been a number of reports on the use of PDMS as an alternative to microfluidic devices for cell experiments (Berthier et al., 2012). In addition to polystyrene (PS) as the conventional material, glass, PMMA, and COP have also been tested. While glass has good optical transparency that is beneficial for observing cells, particularly for fluorescent imaging, it is too fragile for handling and microfabrication with a complicated structure. PS, PMMA, and COP are thermoplastic materials that are applicable for mass production at low costs. Since PS is known to show higher autofluorescence than PMMA and COP, it might interfere with the cellular observations during fluorescence imaging (Young et al., 2013). In the case of PMMA and COP, the solvent bonding process id required for fabrication of the microfluidic devices, and this process might disrupt the microfluidic structure (Ng et al., 2016; Shah et al., 2006) as well as cause leakage from the solvents used, thus causing cellular damage (Wan et al., 2017). PMMA also has issues with generating clacking after alcohol treatments (ethanol or isopropanol) for sterilization prior to use in cell culture. Recently, we established the COP-MPS with a photobonding process instead of solvent bonding to eliminate the drawbacks of the conventional MPS using thermoplastic materials (Yamanaka et al., 2020). This COP-MPS allows culturing of human induced pluripotent stem cells (hiPSCs) (Takahashi et al., 2007; Yu et al., 2007) and reducing undesired apoptotic cells. Thus, our proposed COP-MPS is advantageous for establishing the *in vitro* NAFLD model over the conventional MPS made of glass and other thermoplastic materials with solvent bonding.

In terms of the cell sources, HepaRG cells were used in this study since they are widely used for *in-vitro* NAFLD modeling. However, to obtain physiologically relevant insights, primary human hepatocytes (PHHs) are still the gold standard, but PHHs are difficult to obtain from healthy donors and have limited proliferation. Therefore, other alternatives are utilized, and hiPSCs could be considered as potential candidates owing to their ability for unlimited self-renewal and differentiation to almost any type of tissue. Recently, there have been efforts to apply hiPSC-derived HLCs for pharmaceutical research and disease modeling (Avior et al., 2016; Ramli et al., 2020; Rowe and Daley, 2019; Takayama et al., 2018; Tian et al., 2016). Using hiPSCs, three-dimensional cell culture and organoid generation with better functionalities than those of two-dimensionally cultured hepatocytes (Kamei et al., 2019; Takayama et al., 2018) can be conducted for applications to *in vitro* NAFLD modeling.

Moreover, the symptoms of NAFLD not only manifest in the liver but also in other tissues, such as the intestines and heart. These organs affect each other via the blood stream and thereby affect progression. For example, the gut–liver axis (GLA) is one of the most important components for initiation and progression of NAFLD (Sumida and Yoneda, 2018; Wiest et al., 2017). To elucidate the mechanisms of NAFLD and determine the optimal treatment, such interactions need to be considered. Since the cell lines and primary cells are difficult to obtain from a single donor, observation of the interactions of tissue cells from different donors might cause incorrect results; hiPSCs allow the preparation of multiple tissue cells from the same cell source, and MPS allows recapturing intertissue interactions (Kamei et al., 2017; Trapecar et al., 2020). FFAs act as the mediators of intertissue interactions, and our proposed COP-MPS is expected to be beneficial for observing such interactions without losing the mediators for advancement of *in vitro* NAFLD modeling.

## 5. Conclusion

NAFLD is a critical step in the progression of liver disease, and requires an *in vitro* model to understand the *in vivo* pathological conditions for drug discovery. To establish an *in vitro* NAFLD model, instead of PDMS-MPS, a COP-MPS was proposed and utilized to prevent absorption of FFAs, NAFLD initiators, and AdipoRed fluorescent lipid dye. In light of these results, the COP-MPS has been proved to offer advantages over the PDMS-MPS, especially to establish the *in vitro* NAFLD model. To induce NAFLD-like phenotypes in differentiated HepaRG cells, PA and OA:PA were applied to the COP-MPS, and lipid accumulation was observed in the PA-treated HepaRG cells. To investigate whether the accumulated lipids caused apoptosis, Annexin V apoptotic marker staining was carried out, and it was confirmed that the PA-treated HepaRG cells had the highest rates of apoptotic cells compared to other treatments. Thus, we demonstrated an *in vitro* NAFLD model using COP-MPS. We envision that this model could be applied to drug discovery for the treatment of NAFLD, resulting in reduction of the number of liver transplants and patients with liver diseases.

## Supporting information

Supplemental informations

## Declaration of conflicting interest

M.Y. is an employee of Ushio Inc. A portion of this project was financially supported by Ushio Inc. The other authors declare no competing interests.

## Acknowledgment

Funding was generously provided by the Japan Society for the Promotion of Science (JSPS: 16K14660, and 17H02083) and the LiaoNing Revitalization Talents Program (XLYC1902061). The WPI-iCeMS is supported by the World Premier International Research Center Initiative (WPI), MEXT, Japan.

