## Supplemental informations for "*In vitro* nonalcoholic fatty liver disease model with cyclo-olefin-polymer-based microphysiological systems"

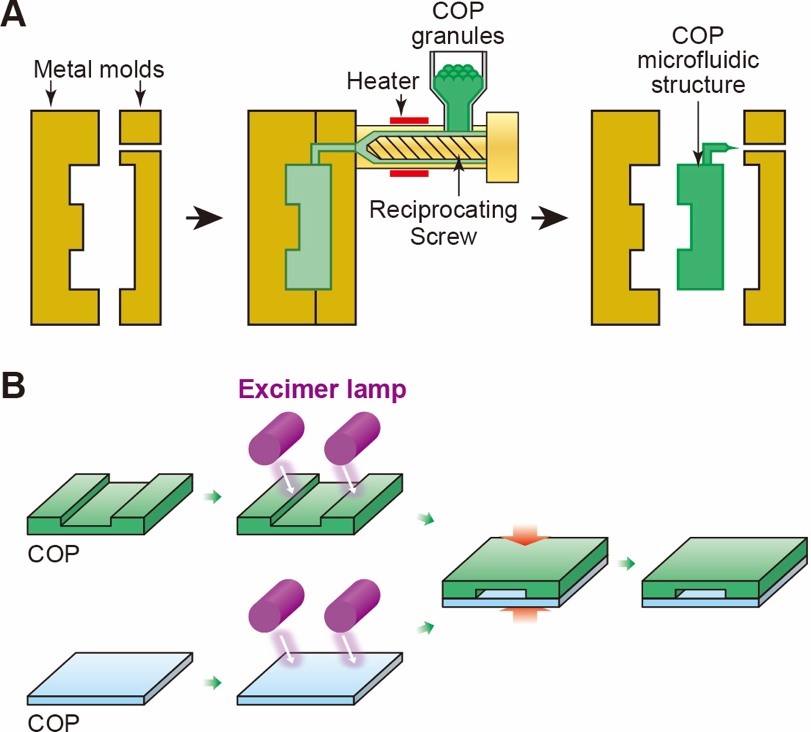

**Fig. S1.** Fabrication of microphysiological system (MPS) made of cyclo-olefin polymer (COP) for modeling nonalcoholic fatty liver disease (NAFLD), namely COP-MPS. (A) Metal molding process to fabricate COP microfluidic structure. (B) Photobonding process to assemble two COP structures using vacuum ultraviolet (VUV, 172 nm) beams from an excimer lamp.

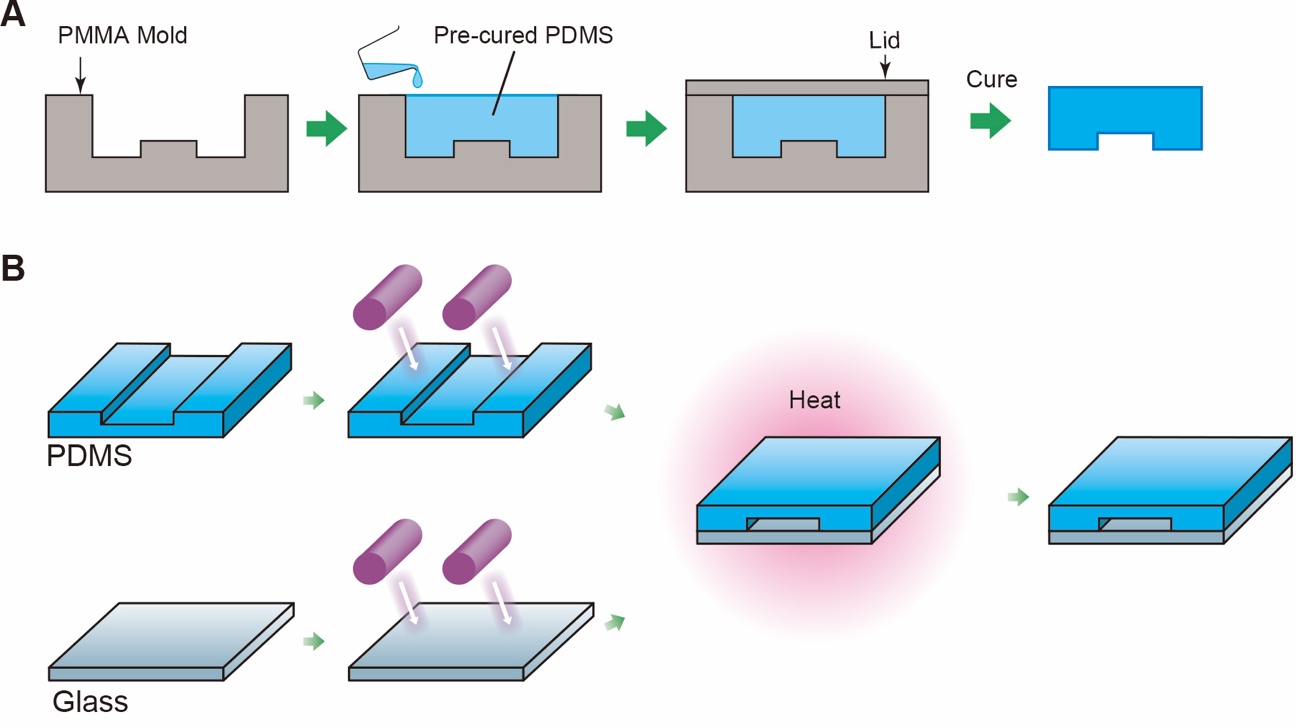

**Fig. S2.** Fabrication of MPS made of polydimethylsiloxane (PDMS) with the same microfluidic structure as COP-MPS, namely PDMS-MPS. (A) Molding process with polymethyl methacrylate (PMMA) mold for the PDMS microfluidic structure. (B) Photobonding process to assemble the PDMS microfluidic structure over a glass slide using VUV from an excimer light source, followed by heat treatment at 80 °C for 24 h.
